# Comparative study of *Anopheles gambiae* larval habitats and larval densities in relation to physicochemical parameters in three urban sites in Abidjan district

**DOI:** 10.1101/2025.04.05.647363

**Authors:** Yves AK Kacou, Firmain N Yokoly, Sandra Le Bissonnais, Julien BZ Zahouli, Jackson IK Kouame, Jean-Philippe B Tia, Benjamin G. Koudou

## Abstract

The knowledge of *Anopheles gambiae* s.l larval habitats is important in order to choose an appropriate malaria control strategy. The purpose of this study is to identify potential mosquito breeding sites in the district of Abidjan and analysed the relationship between environmental factors and larval abundance.

Abiotic characteristics such as temperature, pH, dissolved oxygen, salinity, conductivity and biotic characteristics (larval density) of *Anopheles gambiae* s.l. larval habitats were examined from January to November 2021 in three areas of the Abidjan district. The larval collection was performed using dippers and pipettes.

During the rainy season, the proportion of breeding site was more important in NDotre and in the port area of Treichville in contrast to 43rd BIMA, where furrows predominated. On the other hand, during the dry season, the proportion of breeding sites was higher in 43rd BIMA area, followed by the port area with and NDotre with. The proportions of breeding sites were highly significant between the two seasons in each site. Mean larval densities varied significantly between sites (χ2 = 108.9, df=2, p < 0.001. In Port area and NDotre, temperature was positively correlated with larval density (r=0.2, p < 0.001). In the 43rd BIMA, conductivity (r=0.8, p < 0.001), pH (r=0.64, p < 0.001) and salinity (r=0.2, p < 0.001) were positively correlated with larval density, while dissolved oxygen was negatively correlated in in port area, 43rd BIMA and NDotre (r = −0.01, p < 0.001; r = −0.53, p < 0.001; r = −0.3, p < 0.001) respectively.

This study revealed that the larval habitats of *An. gambiae* s.l. were not very diverse in these areas. These larval habitats were mainly affected by human activity. These data may shed light on malaria control strategies in the cities.

## Introduction

Between 1950 and 2010, the world’s population grew from 2,532 to 6,896 million, an increase of 172%. But within this total, the rural population grew by only 87%, compared with +377% in urban area [1]. While the urbanization rate in Europe and America is around 1%, the fastest urban growth is in Africa, at 4% per year between 2000 and 2020. The natural demographic growth of urban areas, the massive rural exodus from the more or less distant countryside to the cities, the transformation of certain rural localities into cities and the integration of certain rural localities on the outskirts of expanding cities are the reasons behind this accelerated growth of cities [2]. In Sub-Saharan Africa, around 150 million people living in urban areas are exposed to the risk of malaria, and it is estimated that between 6-28% of malaria cases occur in urban areas every year. In Côte d’Ivoire, urbanization is accelerating, with 15,428,957 (52.5%) people currently living in the cities, compared with 13,960,193 (47.5%) in rural areas. The urbanization rate has risen from 32.0% in 1975 to 52.5% in 2021. The city of Abidjan is the most populous in Côte d’Ivoire, with 5,616,633 inhabitants, and is home to 36% of the country’s urban population, with an annual growth rate of 2.9%, i.e. doubling every 24 years, and is the country’s most industrialized city [3]. Unfortunately, this urbanization in sub-Saharan Africa, and particularly in Côte d’Ivoire, is accompanied by the proliferation of vectors for diseases such as malaria [4,5]. Abidjan is divided into two health districts, Abidjan 1 and Abidjan 2, and the incidence of malaria in these two districts is 87,5‰ and 77,08‰ respectively[6]. In Côte d’Ivoire, as in most African countries, malaria vector control relies essentially on the use of long-lasting insecticide-impregnated nets (LLINs) and indoor residual spraying (IRS) [7–9]. Given the resistance of adult mosquitoes to insecticides, malaria control can focus on the larvae of vectors by identifying breeding sites. This becomes an effective alternative in the fight against the disease [10,11]. Larvae control cannot be effective without knowledge of the climatic factors of the larval habitats and the various ecological and physicochemical parameters of the water in which the larvae develop. Abidjan is environmentally heterogeneous, with industrial, market-garden and residential areas. Abidjan presents a heterogeneous environment with the presence of industrial zones, market gardens and residential areas. In Abidjan, dengue vector habitats have been identified in industrial environments [14]. However, the larval habitats and the relationship between abiotic parameters and the abundance of *Anopheles gambiae* larvae, primary vector of malaria in Africa, remain unknown in Abidjan. Larval habitats and physicochemical parameters could vary from one environment to another and certain physicochemical parameters could also influence larval abundance depending on the type of environment.

The aim of the study was to identify the habitats and distribution of *An. gambiae* s.l. larvae in these three environments subject to different pollutants, as well as the physicochemical characteristics of the habitats conducive to larval abundance at these sites. It took place in industrial, vegetable growing and residential environments in the Abidjan district. These data will be used by the National Malaria Control Program to implement strategies to fight the disease.

## Materials and Methods

### Study sites

This study was conducted from January 2021 to November 2021 in the district of Abidjan (Figure 1), which consists of 3 different anthropogenic zones, namely an urban agricultural vegetable growing site (43rd BIMA), an industrial zone (Treichville) and a residential zone (NDotre) (Figure 1).

The 43rd Marine Infantry Battalion (43rd BIMA) (5.265611 N; −3.956709 W) is a neighbourhood located east of Port-Bouet, in the southern part of Abidjan, with a population of 618,795 [3]. The district is home to a market-gardening area along the road to Abidjan’s Félix Houphouet-Boigny international. The market gardening crops grown in this area include lettuce, chili peppers, okra, tomatoes, and eggplant, which have a short cycle (3 to 4 months). The growers use pesticides, fungicides, and other inputs to improve agricultural yields, and some even use non-approved insecticides [15]. The district is located close to the Atlantic Ocean.

The port area of Treichville (5.2811429 N, −4.0087109 W) is located in the commune of Treichville, in the southern part of Abidjan. This commune has a population of 106,552 over an area of 900 hectares [16]. The port area hosts an industrial zone comprising cement, food industry, cosmetics, and pharmaceutical companies. Industrial waste and certain toxic products used for fishing are sources of pollution in this environment. The poor environmental quality exposes the population to vector-borne diseases such as malaria [17].

NDotre (5.4484627N; −4.0823322W) is a sub-district of the commune of Anyama. This commune is located 10 km north of Abidjan and has a population of 389,592 [3]. The residents of NDotre face poor management of household waste and are therefore exposed to environmental diseases such as malaria [18]. The NDotre neighbourhood is home to tens of thousands of inhabitants and has developed rapidly. In addition to the problem of household waste management, the residents of this neighbourhood face inadequate drainage systems and insufficient health infrastructure. The presence of household waste near homes creates an unsanitary environment conducive to the development of disease vectors such as malaria [19]. The choice of these zones considered the degree of urbanization of the sites, the impact of industrial and agricultural activity. All these zones are located in the same geographical region, with the same climate. Abidjan’s climate is sub-equatorial, characterized by two alternating dry and rainy seasons. The long rainy season extends from April to July followed by the short dry season in August. The short rainy season runs from September to November, and the long dry season from December to April. Average annual rainfall is 1,500 mm. The hottest month is February, with an average annual temperature of 27°C and relative humidity is more than 80% [20].

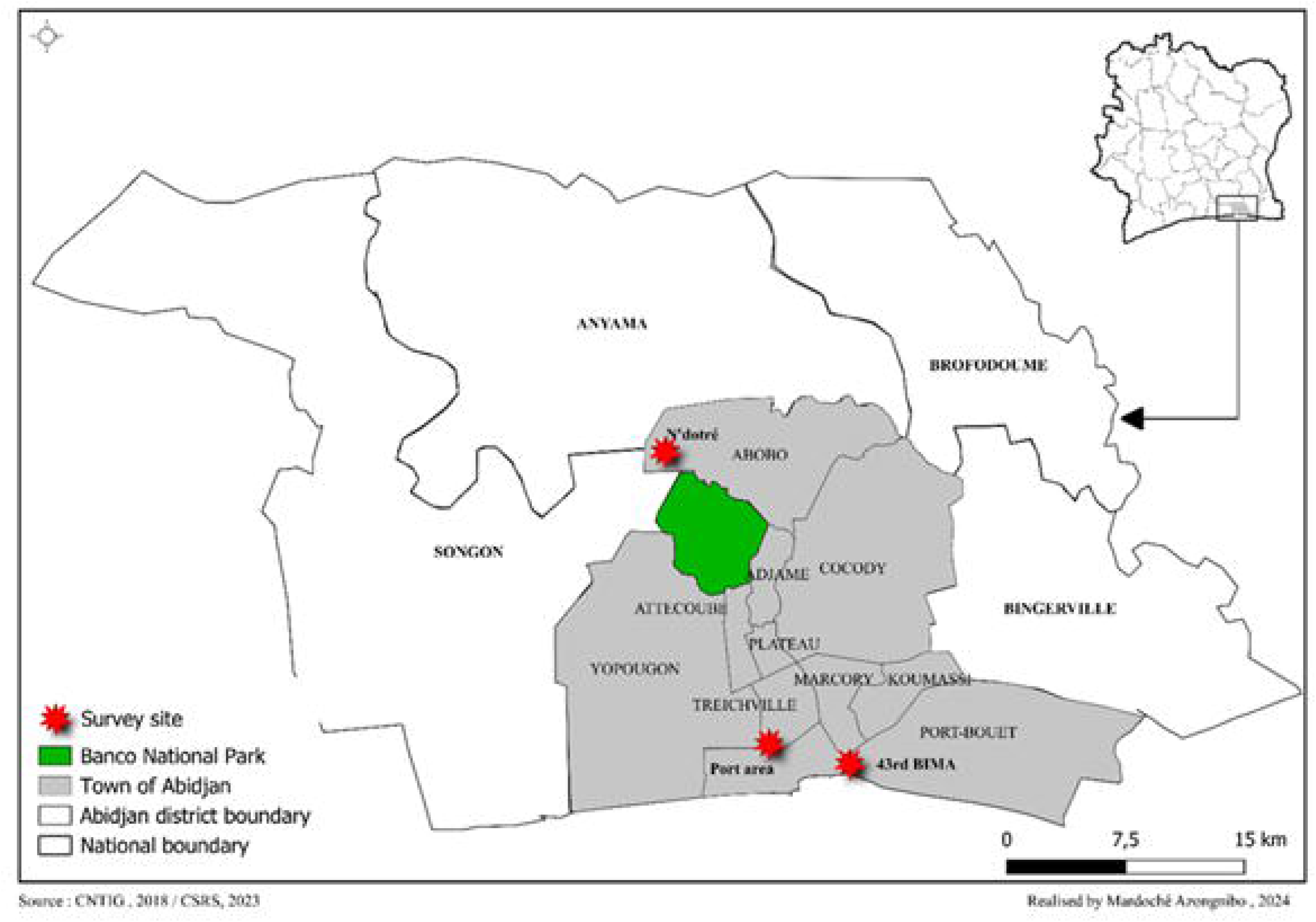

### Larval survey

*Anopheles gambiae* s.l. larvae were collected in industrial, vegetable growing and residential areas. These collection sites are located in the city of Abidjan, notably in the port area of Treichville, the 43rd BIMA and N’Dotré districts. The surveys took place from January to March 2021 and August 2021 during the dry season and from May to July and September to November 2021 during the rainy season. The survey consisted in assessing An. gambiae s.l. breeding sites in these areas. Larval habitat assessments took place over a three-week period in the study areas. Puddles and other objects harbouring Anopheles larvae were studied. Ponds, furrows, gutters, tyres and building materials were considered temporary and artificial habitats. Marshes were natural and permanent habitats. The presence or absence of larvae was determined by visual inspection of breeding habitats. Using the morphological criteria of *Anopheles* larvae at the water’s surface as described by Gillies and Coetzee [21]. Larvae and pupae of *An. gambiae* s.l. were sampled from each positive larval site using a 350 ml dipper, as described by Hessou-Djossou [5].

Ten dipper strokes were made in each larval site, so as to cover the entire surface of the site. Each time, the number of larvae was counted, and the contents poured into plastic containers. The average larval density per larval site was calculated by dividing the number of larvae obtained by the number of dipper strokes. The boxes containing the larvae were labelled and transferred to the CSRS laboratory facilities for rearing.

### Characterization of larval habitats and physicochemical parameters

For each positive habitat, the geographical coordinates were recorded prior to habitat characterization. Larval habitats were characterized by visual observation in the field and classified as, furrows, cistern, marshes, ponds, gutters, abandoned tyres, depending on locality. Physicochemical parameters as salinity (g/L), dissolved oxygen (mg/L), hydrogen potential (pH), conductivity(µS/cm) and temperature were assessed directly in the field during larval surveys, using a HANNA HI 9829 multi-parameter. For each physicochemical parameter, three values were taken, then averaged. This average represented the value of the chemical element sought.

### Static analysis

The statistical analyses were carried out using the R software version 4.1.1 (2021.08.10). The median and interquartile range of the physicochemical parameters of all the habitats has been calculated for each locality and season. The chi-square test was used to compare the proportions of larval habitats by season in each site. However, the Mann-Whitney test was used to compare the medians and interquartile range of the physicochemical parameters, larval density by habitat and the larval density between localities. A principal component analysis (PCA) was used to assess the relationship between physicochemical parameters and larval density by locality. Significance is set at α=0.05 threshold. There is a significant difference if p-value ≤ 0.05.

## Results

### Larval breeding sites

A total of 213 *An. gambiae* s.l breeding sites were surveyed during the rainy and dry seasons at all sites. Overall, the high proportions of breeding sites were found during the rainy season at all sites. In the port area of Treichville, (n = 66, 92.95%) of the breeding sites in the wet season were ponds, compared to (n = 2, 2.81%) tyres and (n = 3, 4.22%) gutters in the dry season. The proportions of breeding sites in the dry season (n = 5, 7.04%) and (n = 66, 92.95%) in the rainy season were highly significant between seasons in the Port area of Treichville (χ² = 52.408; df = 1; p < 0.0001). In 43rd BIMA, only furrows (n = 65, 91.54%) were found during the rainy season. In the dry season, the sites used were marshes (n = 2, 2.81%) and furrows (n = 4, 5.63%). The proportion of breeding sites in the rain season (n = 65, 91.54%) was higher than in the dry season (6, 8.45%). The proportions of breeding sites between seasons were highly significant (χ² = 49.028; df = 1; p <0.0001). Furthermore, in N’Dotré, the proportion of breeding sites in the rain season consisted of ponds (n = 67, 94.36%) compared to (n = 4, 5.63%) (tyres, ponds, cistern) in the dry season. The proportion of breeding sites was highly significant between seasons (χ² = 55.901, df = 1, p <0.0001). In the dry season, breeding sites were consisted of tyres (n=2, 2.81%), followed by ponds (n=1, 1.4%) and cistern (n=1, 1.4%), table 1. In all the sites, a diversity of breeding sites was obtained in the dry season as opposed to the rainy season **(table 1).**

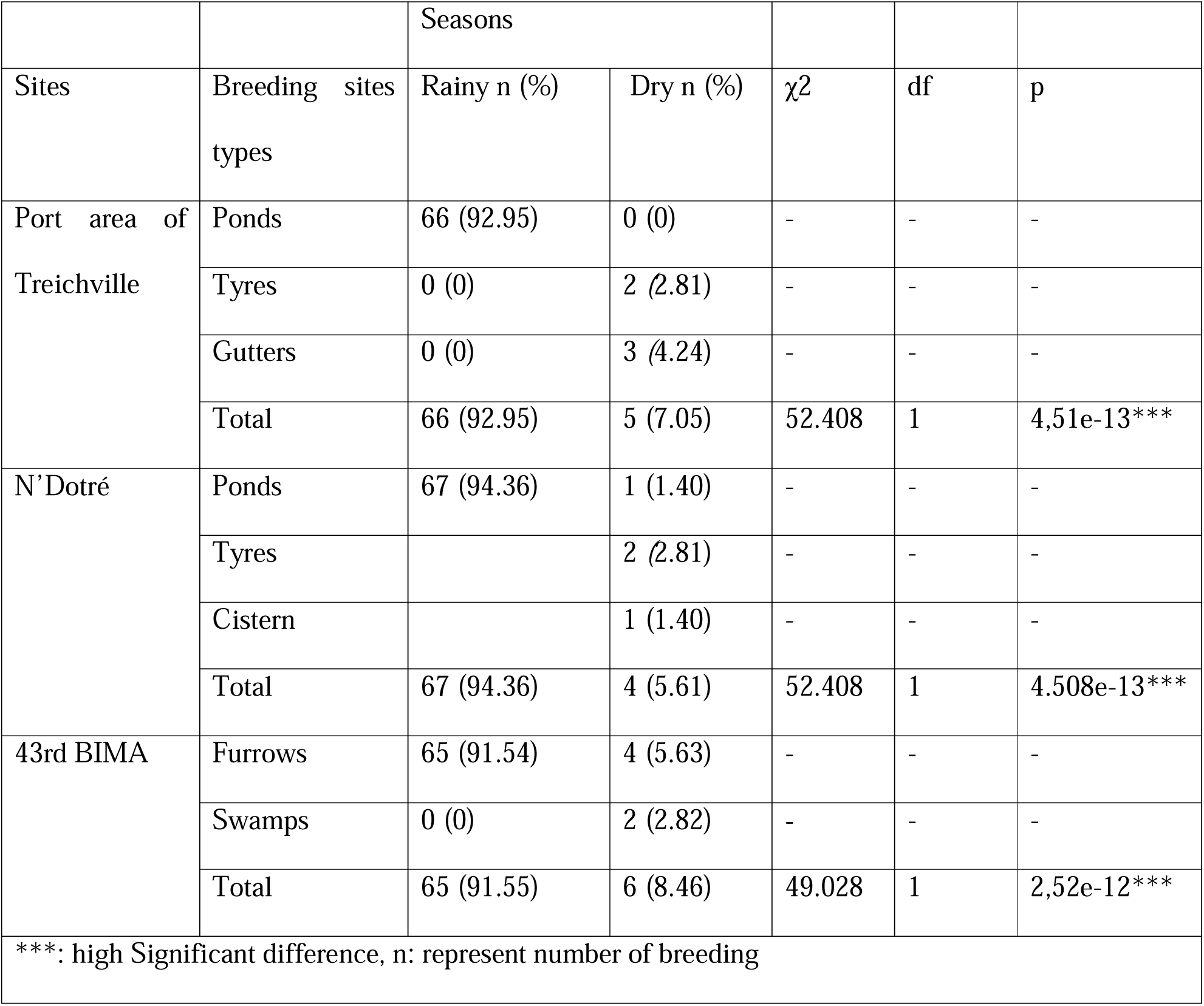

### Productivity of breeding sites

In the Port area of Treichville, larvae were abundant in gutters 28 ± 12 larvae/dipper, then ponds 24.1 ± 8.5 larvae/dipper, and tyres 19 ± 0.5 larvae/dipper. No significant difference was observed in the density of breeding sites (χ²=4.1; df=2; p=0.12). However, in N’Dotré, larval density was higher in ponds 19 ± 6 larvae/dipper, followed by cistern (14 ± NA larvae/dipper). and lower in tyres 1.5 ± 1 larvae/dipper. A significant difference was observed between the densities of breeding sites (χ2=7.2; df=2; p=0.02). In 43rd BIMA, larval densities varied from 12.3±3.8 larvae/dipper in furrows to 10.6±0.53 larvae/dipper in swamps. No significant difference in density between habitats was observed (χ²=0.272; df=1; p=0.06), **(table 2).**

### Larval productivity by season

In the rainy season, the average larval densities were 24.25 ± 8.5 larvae per dip in the Port area, followed by 18.40 ± 5.6 larvae per dip in the N’Dotre, and 12.39 ± 3.8 larvae per dip in 43rd BIMA and were significantly different (χ2=6.152, df = 2, p = 0.04) **(Figure 2).** In the dry season, the average larval densities were 24.08 ± 9.8 larvae per dip, followed by 13.68 ± 17.1 larvae per dip and 11.16 ± 2.1 larvae per dip, respectively in industrial, residential, and agricultural areas (χ2=108.9, df = 2, p<0.001) **(Figure 3).**

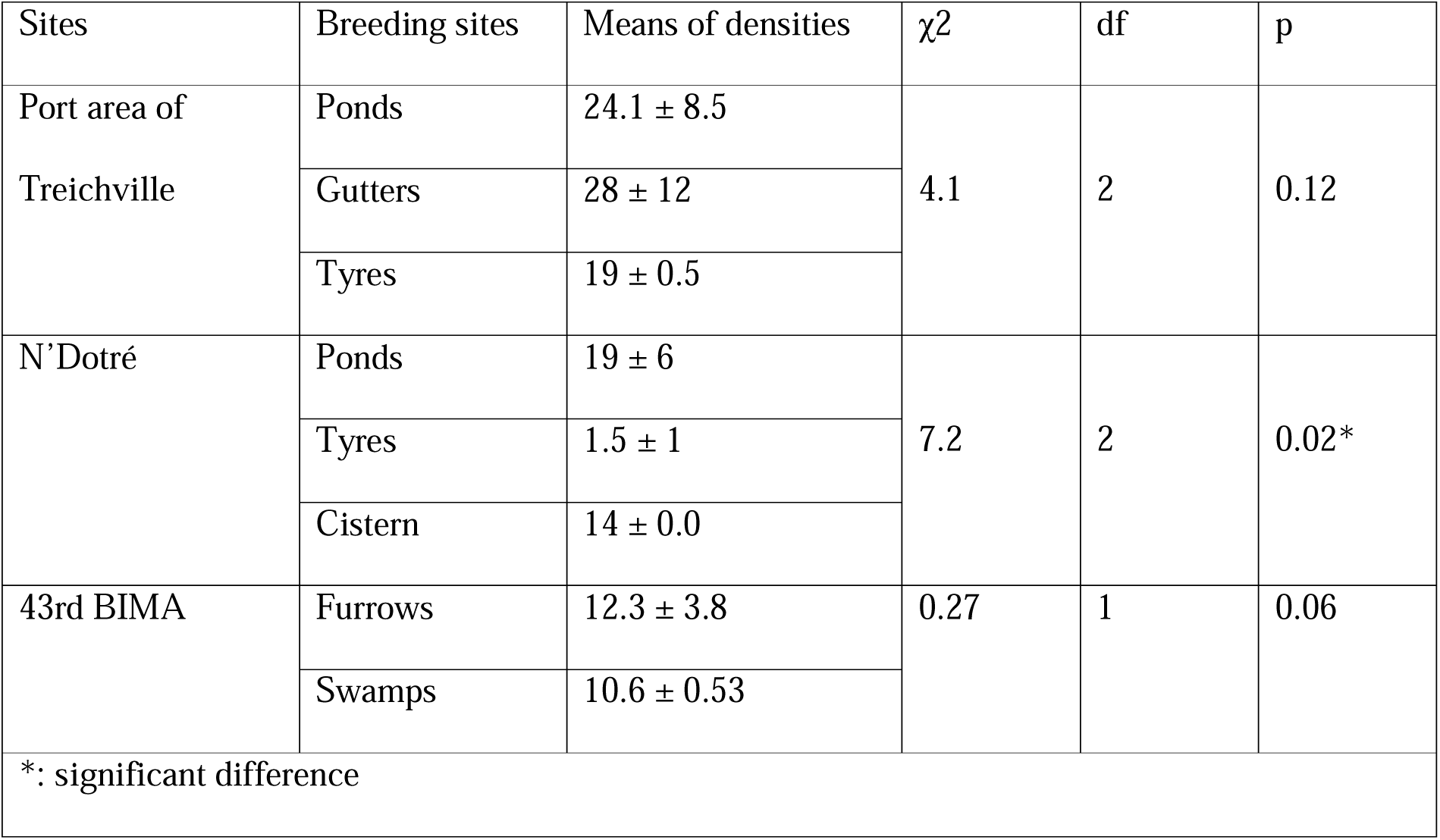

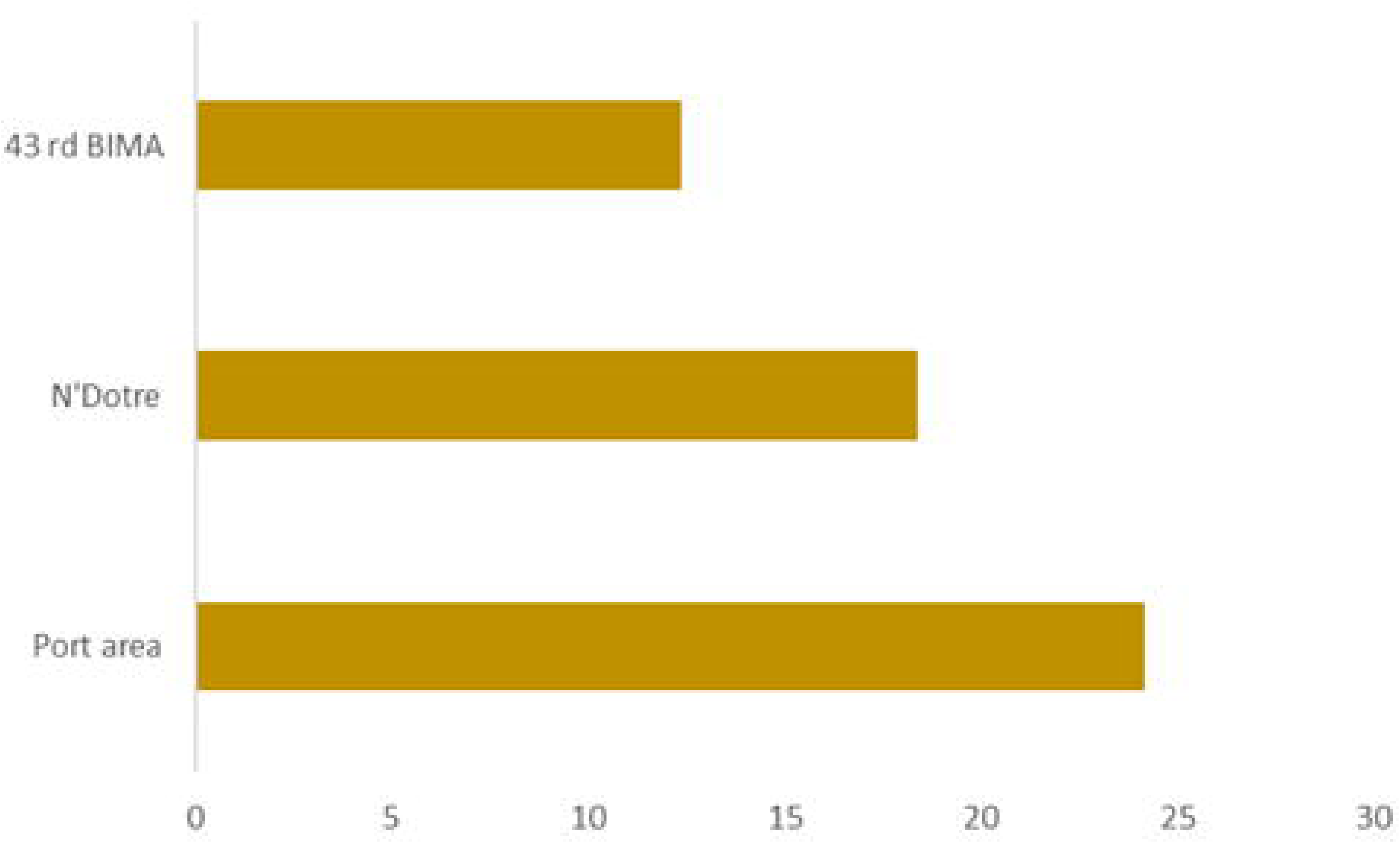

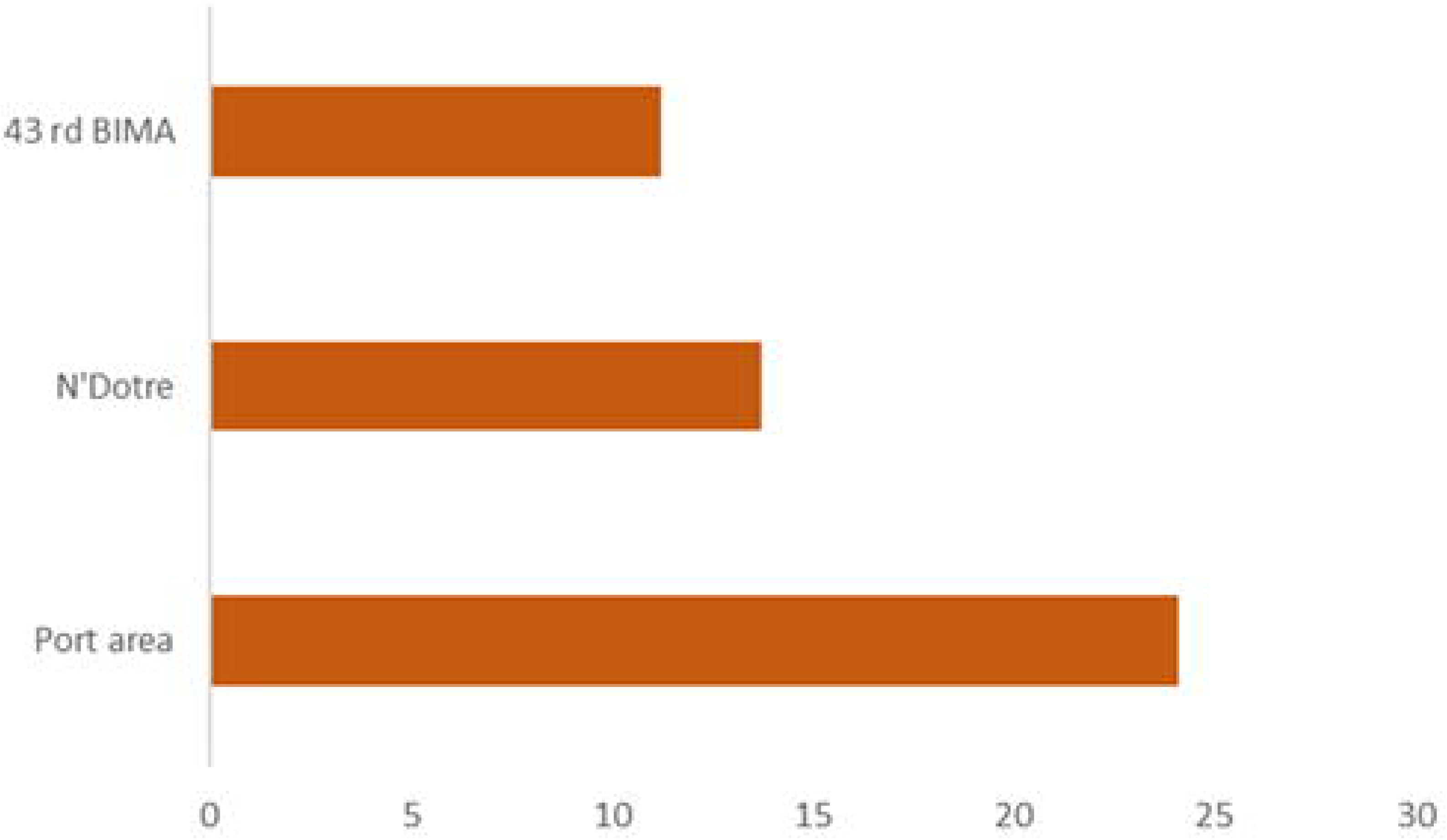

### Seasonal influence on physicochemical parameters

In the rainy season, median and interquartile ranges of dissolved oxygen were 2.4 [1.28-2.74]; 1.7 [0.9-2.34] and 0.4 [0.18-1.8] respectively in the Port area of Treichville, N’Dotré; and 43rd BIMA. These medians were statistically different from one locality to another (χ2= 22.6, df = 2, p<0.001). Medians and interquartile ranges of conductivity were 704 [257-1006] at 43rd BIMA, compared with 434 [300.75-545.50] and 255 [164.9-465] in Port area and N’Dotré respectively. Medians and interquartile of conductivities were different between localities (χ2= 31.5, df = 2, p < 0.001). Medians and interquartile ranges of salinities were 0.2 [0.1-0.5] in 43rd BIMA compared with 0.1 [0.1-0.2] and 0.1 [0-0.2] in Port area and N’Dotré respectively.

Medians and interquartile ranges of salinities were different between localities (χ2 = 20.6, df = 2, p < 0.001).

Median and interquartile ranges of pH at were 9.3 [8.9-9.9] in Port area and N’Dotré respectively, and 8.2 [7.5-8.63] in 43rd BIMA, pH medians and interquartile ranges differed within localities (χ2=66.7, df = 2, p < 0.001). Median and interquartile ranges of temperatures were 32.6 [30.7-34] in 43rd BIMA, 32 [30.1-33.4] in N’Dotré, and 29.5 [28.6-34.8] in Port area. Median and interquartile ranges of temperatures were significantly different between localities (χ2=9.8, df = 2, p= 0.007).

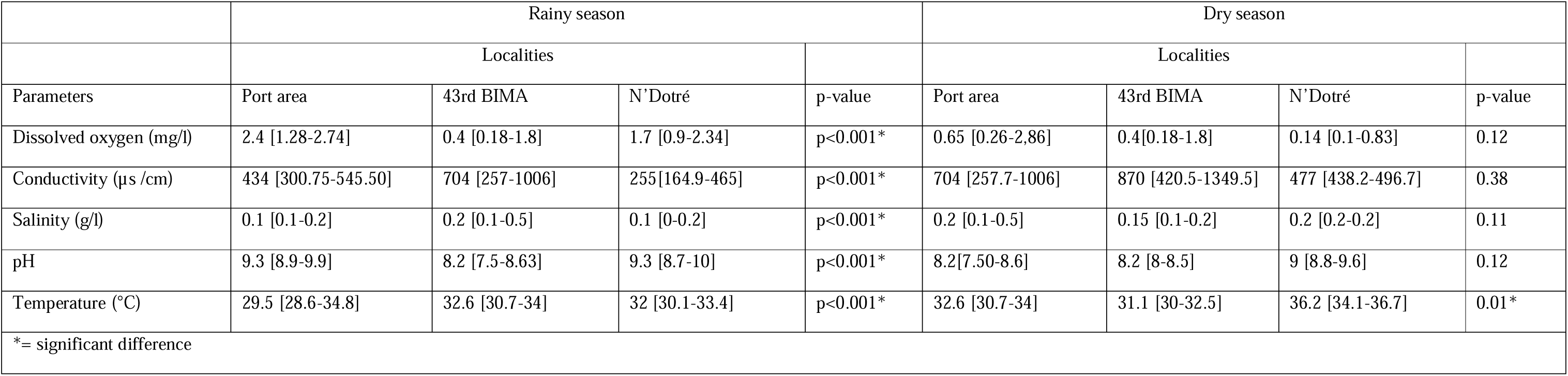

In the dry season, median and interquartile ranges of dissolved oxygen were 0.65 [0.26-2,86]; 0.4[0.18-1.8] and 0.14 [0.1-0.83] at the Port area of Treichville, 43rd BIMA and N’Dotré respectively. No significant difference was observed for this parameter (χ2=3.4, df= 2, p = 0.17). The median and interquartile ranges of conductivities were 870 [420.5-1349.5] at 43rd BIMA, compared with 704 [257.7-1006] and 477 [438.2-496.7] at Port area and N’Dotré respectively. The median and interquartile ranges of conductivities were statistically similar (χ2= 1.9, df = 2, p = 0.38).

Medians and interquartile ranges of salinities were 0.2 [0.2-0.2] in NDotre compared with 0.2 [0.1-0.5] and 0.15 [0.1-0.2] in Port area and 43rd BIMA, respectively. Medians and interquartile ranges of salinities were not different between localities (χ2 = 2.1, df = 2, p = 0.11). Median and interquartile ranges of pH at were 9 [8.8-9.6] in N’Dotré, 8.2 [8-8.5] and 8.2[7.50-8.6] in 43rd BIMA and Port area respectively, pH medians and interquartile ranges not differed within localities (χ2 = 4.2, df = 2, p = 0.12). Median and interquartile ranges of temperature at N’Dotré was 36.2 [34.1-36.7], while Port area and 43rd BIMA, the median and interquartile ranges temperatures were 32.6 [30.7-34] and 31.1 [30-32.5]. The medians and interquartile of temperatures were significantly different (χ2=8.7, df = 2, p = 0.01). **(table 3).**

### Principal component analyses

Principal component analysis showed correlations between physicochemical variables and larval density in each zones. In the three study sites, dimensions 1 and 2 explain all the variables in the table 4.

In the Port area of Treichville, the data represented on dimensions 1 and 2 explained 57.5% of the total variable. In these two localities, pH and temperature were weakly associated with dimension 1 (coefficients r =0.4, r= 0.2). Of these two parameters, pH had the greatest influence on larval density in the Port area industrial zone (r=0.4, p<0.001). Salinity and conductivity were weakly associated with dimension 1 (coefficients r = −0.14, p = 4.08e-16; r = −0.2, p < 0.001), while dissolved oxygen was negatively associated with dimension 1. Dissolved oxygen had a negative influence on the larval density of *Anopheles* populations in the industrial environment of Port area (coeffiecient r = −0.01, p < 0.001) **(Figure 4 a).**

|  |  | Dimension 1 | Dimension 2 | Dimension 3 | Dimension 4 | Dimension 5 |
| --- | --- | --- | --- | --- | --- | --- |
| Sites |  | r | r | r | r | r |
| Port area | Dissolved oxygen | -0.01 | -0.63 | 0.31 | -0.26 | 0.65 |
|  | Conductivity | -0.20 | 0.83 | 0.02 | 0.002 | 0.18 |
|  | pH | 0.40 | -0.17 | 0.36 | 0.81 | 0.06 |
|  | Salinity | -0.14 | 0.78 | 0.24 | 0.07 | 0.35 |
|  | Temperature | 0.20 | -0.03 | -0.87 | 0.24 | 0.36 |
|  | Larvae number | 0.96 | 0.18 | 0.04 | -0.19 | -0.002 |
|  | Density | 0.96 | 0.18 | 0.040 | -0.19 | -0.002 |
| 43rd BIMA | Dissolved oxygen | -0.53 | 0.21 | 0.53 | 0.58 | -0.10 |
|  | Conductivity | 0.80 | -0.27 | 0.07 | 0.01 | 0.36 |
|  | pH | 0.64 | -0.48 | 0.21 | 0.38 | 0.19 |
|  | Salinity | 0.21 | -0.04 | 0.86 | -0.44 | -0.09 |
|  | Temperature | -0.50 | 0.62 | 0.12 | -0.07 | 0.56 |
|  | Larvae number | 0.74 | 0.65 | -0.02 | 0.06 | -0.11 |
|  | Density | 0.74 | 0.65 | -0.02 | 0.06 | -0.11 |
| N'Dotré | Dissolved oxygen | -0.302 | 0.37 | -0.48 | 0.58 | 0.41 |
|  | Conductivity | -0.0080 | 0.72 | 0.51 | 0.14 | -0.09 |
|  | pH | 0.4 | -0.47 | 0.34 | 0.1 | 0.67 |
|  | Salinity | 0.05 | 0.75 | 0.40 | -0.19 | 0.23 |
|  | Temperature | 0.1 | -0.42 | 0.58 | 0.61 | -0.18 |
|  | Larvae number | 0.95 | 0.18 | -0.21 | 0.06 | -0.06 |
|  | Density | 0.95 | 0.18 | -0.21 | 0.06 | -0.06 |
| r: correlation coefficient |  |  |  |  |  |  |

At farming vegetable zone in 43rd BIMA, dimensions 1 and 2 accounted for the total variable Conductivity and pH were strongly correlated with dimension 1 (r = 0.80; r = 0.64).Conductivity and pH were the abiotic parameters that contributed to larval abundance in the 43rd BIMA farming vegetable environment (r = 0.80, p < 0.001; r = 0.64, p < 0.001). Dissolved oxygen and temperature were negatively associated with dimension 1 (r = −0.53, r = −0.50). These two parameters had a negative influence on the larval density of *Anopheles* populations in the 43rd BIMA farming vegetable zone (r = −0.53, p < 0.001; r= −0.50, p < 0.001) **(Figure 4 b).** In the residential environment of N’Dotré, the data represented on dimensions 1 and 2 explained the total variable. In this localitie, pH and temperature were weakly associated with dimension 1 (coefficients r = 0.40, r = 0.1). Of these two parameters, pH had a greater influence on larval density in the residential area of N’Dotré (r = 0.4, p < 0.001). Dissolved oxygen was negatively associated with dimension 1, and negatively influenced the larval density of *Anopheles* populations in N’Dotré (r = −0.30, p = 1.04e-02). Salinity was weakly associated with dimension 1 (r=0.05; p < 0.001) and conductivity was negatively associated with dimension 1 (coefficients r= −0.008; p < 0.001) **(Figure 4 c).**

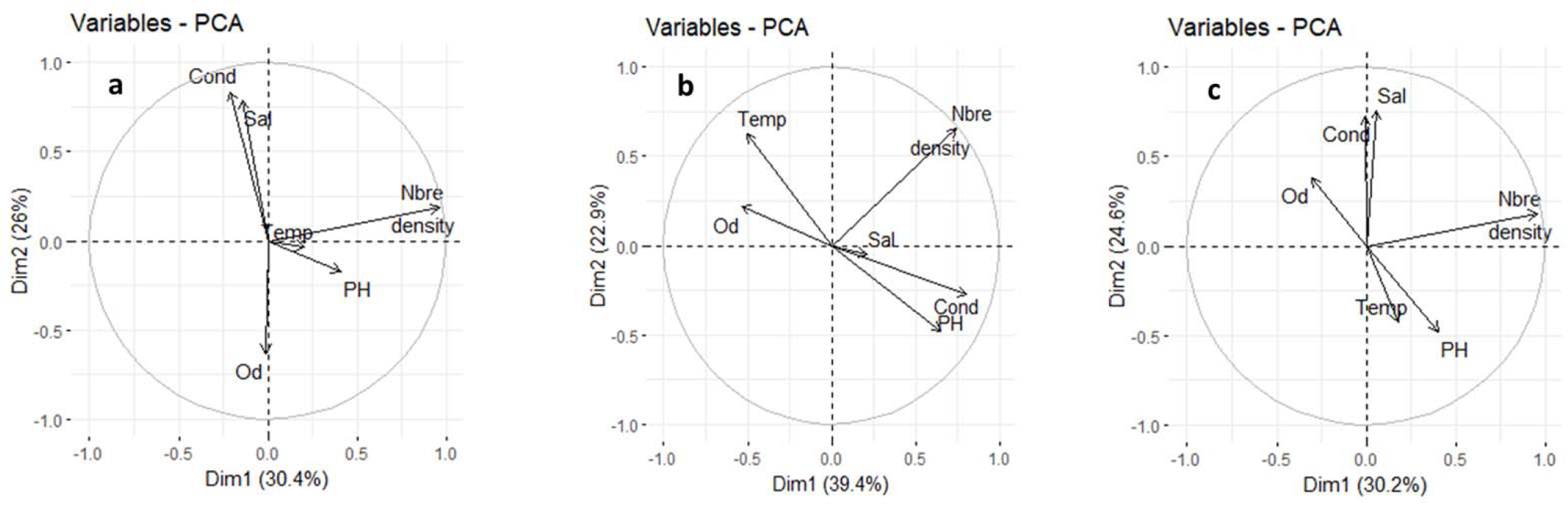

## Discussion

This study was carried out to identify the larval habitats and physicochemical parameters contributing to the proliferation of *An. gambiae* s.l. larvae in industrial, vegetable growing and residential settings in the district of Abidjan, and largest urbanization area of Côte d’Ivoire. Furrows and swamps were the main habitats encountered in urban farming vegetable areas in 43rd BIMA, while in industrial settings at the Port area of Treichville and residential area in NDotre, the habitats were ponds, tyres, gutters and cisterns. These results show that in these areas, the larval habitats of *An. gambiae* s.l. are diversified. The diversity of breeding sites in the study sites could be justified by the urban nature of study sites.

In the industrial zone of Port area, poor waste management and inadequate drainage systems are thought to be the causes of the diversity of larval habitats. In NDotre (residential area), uncontrolled urbanization and the lack of sanitation systems such as sewers are thought to be the causes of the proliferation of larval habitats. In 43rd BIMA (market gardening area), poor management of the water used to water vegetables is thought to be the cause of the proliferation of larval habitats. Our results are in line with trends observed on the diversity of *An. Gambiae* s.l. breeding sites in African and Ivorian urban environments [5,12,22]. On the other hand, the presence of larval breeding sites observed could be due to the intensification of industrial, urban vegetable growing and residential activities, as well as to inadequate sanitation infrastructures in these areas, as previously indicated by Silue [14]. This inadequacy of drainage systems is due to uncontrolled urbanization, as is the case in most of the Sub-Saharan African cities [4]. The presence of *An. gambiae* s.l. in atypical sites (tyres, cisterns, gutters) in industrial and residential areas is due to poor management of these objects in these two environments. *Anopheles gambiae* s.l. had been observed several times in these materials in urban environments [4,23,24]. Atypical breeding sites were only observed in the dry season in residential and industrial environments. The presence of *An. gambiae* s.l. in atypical sites may be linked on one hand, to the lack of pools of water in the dry season and, on the other hand, to the adaptive capacity of *An. gambiae* s.l. to the urban environment. Several publications have reported the presence of *An. gambiae* s.l. in atypical habitats elsewhere and in Côte d’Ivoire [12,14]. This is a good opportunity to draw people’s attention to the need for proper management of these devices to avoid malaria epidemics in the district of Abidjan, as is currently the case with the dengue epidemic observed in the same area [25]. Significant differences were observed between physicochemical parameters in the rainy season. This underlines the importance of rain in larval abundance. Average dissolved oxygen is low and varies according to the environment, so it would indicate pollution of larval habitats. Soil contamination by industrial, agricultural and household pollutants in industrial, vegetable growing, and residential areas respectively would explain the low dissolved oxygen values in the various localities. Despite the presence of pollutants of various origins, the larvae thrive. Yet this species is renowned for its preference for clear, unpolluted water. *Anopheles* larvae have been observed in polluted urban waters in Côte d’Ivoire and Cameroon [26,27]. The highest average salinity in this study was 0.4 g/l or 400 parts per million (ppm). In Cameroon, the mean salinity value in larval habitats of *An. gambiae* s.l. was 97.6 (ppm) in urban environment. Of these two values, it would seem that the mean salinities obtained in this study was very high compared to those obtained in Kribi (Cameroun) [28]. The salinity values obtained in this study would indicate the low brackish nature of the larval habitats surveyed. *Anopheles gambiae* s.l. is known for its preference for slightly brackish water [29,30]. The highest temperature was 34.67°C in the residential area in the dry season and the lowest was 31.23°C in the industrial environment. The temperatures obtained in our study would be conducive to larval development in our various study sites. The temperature values obtained in this study are consistent with the observations of Huang And Collaborators [31]. Huang *et al*. showed that water and mud temperatures in larval habitats rarely exceeded 35°C [31]. These temperatures would be necessary for the development of the microorganisms that feed *Anopheles* larvae [32,33]. The highest average pH was found in industrial environments (9.57), compared with 9.38 and 8.13 in residential and market garden environments respectively. These results show that the larvae would have developed in alkaline environments. Our results differ from those of Ondo state north in Nigeria. The author have been reported the development of *Anopheles* larvae in acidic environments [34]. This alkalization is linked to the activities of the various microorganisms present in the larval environment. Indeed, the use of urea as an input in vegetable growing activities and the accumulation of human waste in industrial and residential environments would facilitate the ammonification reaction. Under the influence of microorganisms in different larval environments, the ammonification reaction would lead to alkalization of larval habitats [35].

The density of larvae was higher in the port area than in the other two areas in both seasons. This high density of larvae could be explained, on the one hand, by the presence of the port, which offers day and night activities, thus generating a permanent human presence in the area, and, on the other hand, by the availability of larval breeding sites in the area. These two conditions are necessary for the proliferation of *Anopheles gambiae* larvae [36]. In the vegetable-growing zone, the unreasonable use of insecticides could be a factor in the low larval density. Insecticides reduce the fecundity of female *Anopheles*, which could affect their egg-laying capacity, as shown by a study in Tanzania [37]. In addition to reducing fecundity, the application of certain insecticides such as pyrethroids would act as repellents for *Anopheles*. This repulsion of the *Anopheles* could lead them to lay their eggs in other insecticide-free areas such as marshes, hence the low larval density [38]. The frequency of insecticide applications could be one of the causes of low larval density in vegetable growing zone. The correlation between physicochemical parameters and larval density showed a positive correlation between density and conductivity in urban vegetable-growing area. The average conductivity is high, reflecting rapid degradation of organic matter. The high conductivity is linked to the use of fertilizers in this area which could be a source of food for the larvae. This corroborates observations made in East Africa and Ghana, where larval density was positively correlated with conductivity following fertilizer and pesticide applications [39,40]. In industrial and residential areas, larval density is positively correlated with temperature. This would suggest that temperature is an important factor in larval proliferation. In fact, temperature not only activates the activity of microorganisms in the soil, but also produces more available food for the larvae. Temperature has several times been recognized as a determining factor in the development process of *Anopheles* larvae [41,42]. In addition to temperature, pH is also a variable involved in the proliferation of larvae in residential and industrial environments. Temperature and pH act synergistically to promote the proliferation of *Anopheles* larvae. Indeed, temperature would be favourable to the enzymatic activity of certain microorganisms for the degradation of domestic waste, and the degradation of domestic waste could lead to alkalization of larval habitats through the ammonification process. This result is contrary to obtained in Algeria, where *An. gambiae* s.l. larval density was correlated with acid pH [41]. Dissolved oxygen was negatively correlated with larval density in all study areas. In previous studies, a positive correlation was found between *Anopheles* larval density and dissolved oxygen. Certainly, the dissolved oxygen values obtained in this study would be threshold values for *Anopheles* larvae proliferation. Moreover, this species is known to prefer oxygenated habitats [43]. Dissolved oxygen could therefore be a limiting factor for the proliferation of *Anopheles* larvae in this study. It would be even more interesting to conduct studies to find threshold values of dissolved oxygen below which *An. gambiae* s.l. cannot develop. In this study, we show the habitats of *An. gambiae* s.l. could be considered polluted. This pollution of habitats would be due on the one hand to the presence of industrial, domestic, and agricultural pollutants in the habitats, and on the other hand to the urban nature of the study sites. *Anopheles gambiae* s.l. was observed in polluted urban sites prior to our study [44]. While conductivity, pH and temperature had a positive influence on larval densities in vegetable growing, industrial and residential areas respectively, salinity also played an important role in larval proliferation. Indeed, low-salinity habitats are the preferred habitats of *An. gambiae* s.l., as was the case in this study. Low salt levels prevent the development of certain insects that predate Anopheles larvae. Predatory organisms such as dragonfly nymphs and bugs are known to feed on mosquito larvae. Their predation can considerably reduce the density of *An. gambiae* s.l. larvae in larval habitats and considerably reduce the number of adult mosquitoes [45]. However, larvae of this species have been found in habitats with higher salt levels than ours [46,47]. Although this study has shown the relationship between physicochemical parameters and larval density in each environment, it remains limited in identifying microorganisms and their relationship with larval density in each habitat category. Further analysis to identify chemical pollutants that may impact larval development and mapping of larval habitats are also needed to counter the development of *An. gambiae* s.l. larvae in Abidjan city.

## Conclusion

This study revealed a diversity of *An. gambiae* larval habitats generated in three different anthropogenic environments (industrial, market gardening and residential activities) in Abidjan. In these three environments, the larval density varied from one zone to another with larval abundance in the industrial environment. In addition, physicochemical parameters such as pH and temperature were correlated with the abundance of *Anopheles* larvae. While dissolved oxygen had a negative influence on larval development. In view of the results of this study, new malaria control measures such as mapping larval habitats and sanitation measures should be prioritized by the national malaria control program in these areas and throughout the entire Abidjan district.

## Acknowledgements

We would like to express our gratitude to all those who have contributed to the progress of this work in any way. Many thanks to the Centre Suisse de Recherche Scientifique (CSRS) in Côte d’Ivoire. Thanks also to the laboratory technicians of the Swiss Centre and particularly to Mr. Assamoi Jean-Baptiste for their technical support to the laboratory. I could not finish without thanking the Doctors and PhD students for their moral support.

## Authors’ contribution

YAKK: Conceptualization, Methodology, Investigation, Formal analysis, Data curation, Visualization, Writing - original draft, Writing - review & editing CE: supervision and drafted the manuscript, Resources, FNY: drafted the manuscript, JC: reviewed the manuscript for improvement, JBZZ: Writing - original draft, Writing - review & editing, SLB: Writing - review & editing, CE, JKIK: Data analysis, drafted the manuscript, JPTB: Review & editing, BGK: Project administration Methodology, Writing – review & editing, Validation, Supervision.

## Competing Interests

No competing interests exist.

## Notes

### Competing Interest Statement

The authors have declared no competing interest.

